# MagnoliidsGDB: An integrated functional genomics database for Magnoliids

**DOI:** 10.1101/2024.08.19.608005

**Authors:** Yayu Chen, Zhuang Yang, Jiajie Chen, Ping Li, Xinglong Zhao, Sihui Huang, Zhumao Li, Sishu Huang, Jie Luo, Haiyan Hu, Yuanhao Ding

## Abstract

In this study, we present MagnoliidsGDB, an integrated functional genomics database specifically developed for magnoliids, a group of plants known for their unique characteristics and medicinal properties. As of March 2024, MagnoliidsGDB has collected 31 genome sequences from 24 species, 149 resequencing datastes from 4 species, 842 transcriptomic datasets from 20 species, and metabolomic data from 15 species. This comprehensive resource allows researchers to explore the genomic, transcriptomic, and metabolomic variations of magnoliids. The user-friendly search and analysis functionalities of MagnoliidsGDB facilitate the study of genomic variations, gene functions, plant evolution and key metabolites of magnoliids. We hope this database will contribute to the study of magnoliids and increase our understanding of the origin, variation, and molecular basis of the unique biological features of magnoliids.

---

Charles Darwin’s “Abominable Mystery” about the sudden appearance and diversification of dicotyledons in the Cretaceous has puzzled scientists for over a century (Zuntini et al., 2024). As the third largest group of angiosperms (Wu et al., 2021), magnoliids are dicotyledonous plants that retain some primitive anatomic and morphological characteristics, such as single aperture pollen, unfused carpels, small embryos and copious endosperm seeds (James W. Byng, 2016). These features give magnoliids pivotal roles in evolutionary status and may be key to unveiling Darwin’s “Abominable Mystery”. However, the taxonomic position between magnoliids and other angiosperms (especially monocots and eudicots) remains uncertain, and related debates continue (Jiang et al., 2024; Talavera et al., 2023).

Magnoliids comprise four orders: Laurales, Canellales, Piperales, and Magnoliales, with 18 families and over 9, 000 species (Shen et al., 2021). Many species within this group possess specialized biological characteristics and metabolites with strong bioactivities, making them valuable genetic resources and sources of important bioactive compounds for modern medicine. Since the first publication of the genome of magnoliids, *Liriodendron chinense* in 2019 (Chen et al., 2019), a total of 23 species have been sequenced up to March 2024 (Figure 1A). An increasing number of researchers are engaging in related studies, and massive amounts of transcriptomic, genome resequencing, and metabolic data are being rapidly produced (Qin et al., 2021). It is well known that an integrated database can greatly facilitate foundational research on related plants and fields (Li et al., 2023; Yang et al., 2023). However, a reliable platform for the quick accession and utilization of magnoliids genomic information remains unavailable. A genomic database of magnoliids is tremendously useful for researchers not only to study the origin of dicotyledons but also the molecular basis of the unique biological characteristics of magnoliids.

**Figure 1.**
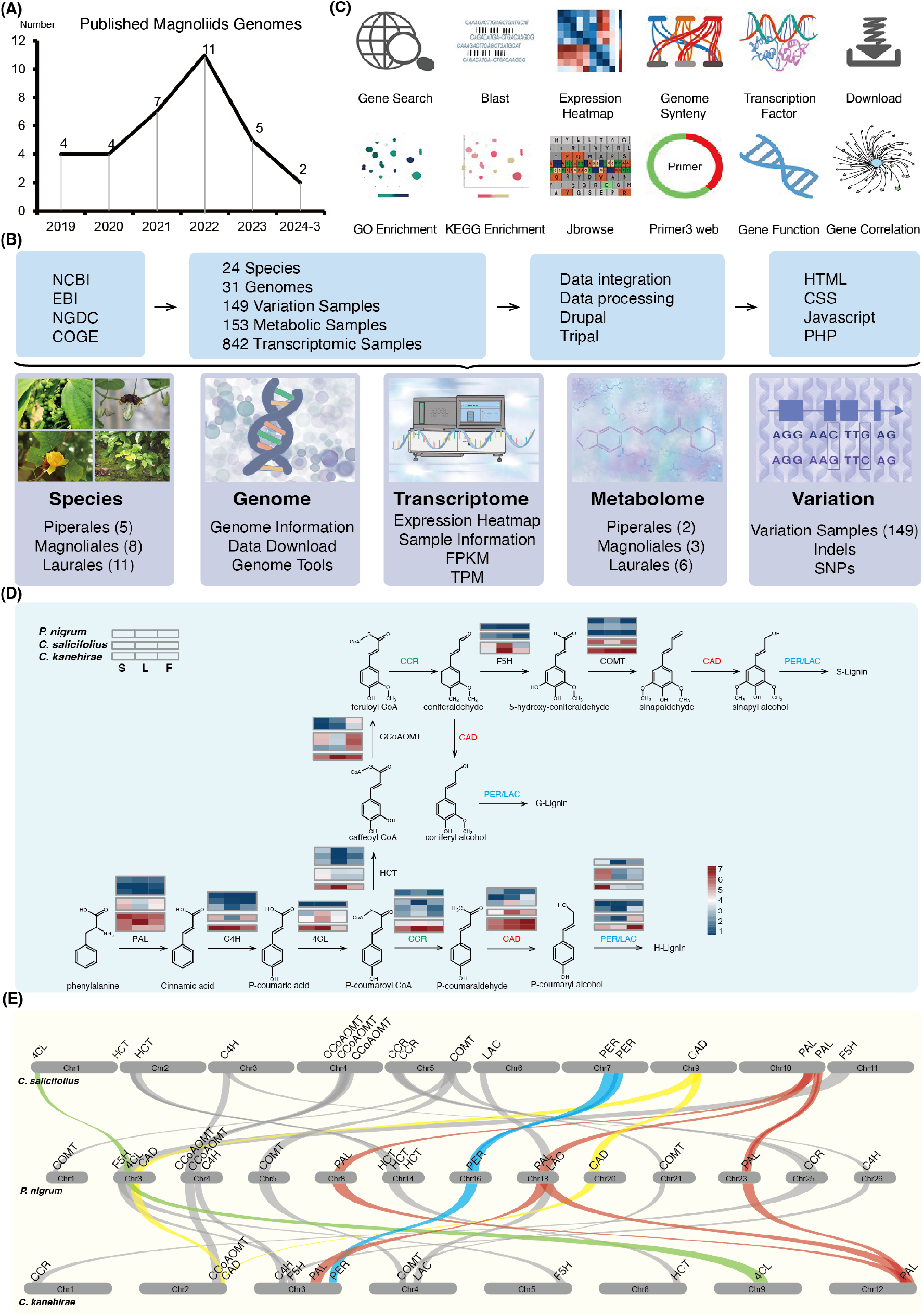
Overview of MagnoliidsGDB structure and functions, alongside an illustrative study of lignin biosynthetic differences in magnoliids. **(A)** Published genomes within the magnoliids clade from 2019 to March 2024. **(B)** The origins of website data, a summary of data information, the steps and programming languages employed in website construction, as well as an overview of the principal website modules. **(C)** An overview of the tools available in MagnoliidsGDB. **(D)** Heatmap tools showing the expression changes of genes from lignin biosynthesis pathway in the stems, leaves and flowers of *Piper nigrum, Chimonanthus salicifolius*, and *Cinnamomum kanehirae*. **(E)** The chromosomal localization of genes related to lignin biosynthesis across three magnoliids species.

Here, we present an integrated multifunctional Magnoliids Genomics Database, MagnoliidsGDB (http://www.magnoliadb.com:7777/), providing abundant multi-omics resources and convenient tools to enable researchers to unravel the evolutionary history, gene functions, and species-special features of magnoliids. As of March 2024, MagnoliidsGDB has collected 31 genome sequences from 24 species, 149 resequencing data from four species, 842 transcriptomic data from 20 species, and metabolomic data from 15 species. The metabolomic data for *Piper nigrum* (Lv et al., 2024), *Piper longum*, and *Persea americana* were obtained through LC-MS-based metabolic analysis (Figure 1B) and provides 11 convenient tools for users to search and analyse (Figure 1C). Researchers can use this platform to obtain information and data on magnoliids conveniently and quickly. MagnoliidsGDB supports multiple analyses of biological function, and provides an upload interface for original data, allowing researchers to efficiently and comprehensively obtain standardized data on magnoliids, thereby promoting and accelerating related research.

The database of magnoliids is constructed with five main modules, (1) Species, (2) Genome, (3) Transcriptome, (4) Metabolome, and (5) Variation (Figure 1B). The “Species” module presents basic information on 24 sequenced magnoliids plants, including chromosome number, common name, distribution, detailed description, image for each species, cross-reference, publication information, and usable hyperlinks to other website or modules. Clicking the “cross-reference” can jump to the NCBI Taxonomy Browser database and PubMed database conveniently to obtain more species information and publications of relevant articles.

The “Genome” module contains 31 genomes, including their published journals, publish date, sequence method, assembly method, genome size, genome coverage, scaffold N50, contig N50, etc. FASTA-formatted files for assembled genomes, CDS sequences, Gff3 file, and protein sequences can be downloaded from this module. When the user clicks on the secondary menu “Jbrowse”, it automatically jumps to the genome browser interface. The module also provides tools related to genome analysis, which can be accessed directly by clicking to navigate to the specified page.

The “Transcriptome” module contains 842 samples from 89 different bioprojects of 20 species (*Aristolochia contorta* (18), *Aristolochia fimbriata* (34), *Piper nigrum* (47), *Piper longum* (3), *Saururus chinensis* (7), *Annona glabra* (1), *Liriodendron chinense* (125), *Michelia alba* (21), *Magnolia biondii* (6), *Magnolia officinalis* (4), *Chimonanthus praecox* (104), *Chimonanthus salicifolius* (30), *Cinnamomum burmannii* (7), *Cinnamomum camphora* (73), *Cinnamomum kanehirae* (8), *Lindera glauca* (21), *Lindera megaphylla* (21), *Litsea cubeba* (87), *Persea americana* (159), *Phoebe bournei* (66)) (Table S1), which have been processed according to the unified transcriptomic analyses methods, and gene expression profiles, including FPKM files and TPM files, can be download from this module. Additionally, the module also provides essential transcriptomic information for users, such as sample names, BioProject, SRR, tissue, stage, age, and country, along with project numbers of samples. Furthermore, we offer users the functionality of gene expression level heatmaps. Users can select their interested species, genes, and analyses to generate corresponding heatmaps. They can also choose the arrangement of the heatmap based on tissue, development stage, biomaterial, accession or project.

The “Metabolome” module comprises metabolic information from 153 samples of 15 species (Table S2), including quantitative and qualitative data, as well as sample information. Upon clicking the name of a metabolite, users are automatically redirected to the corresponding compound’s location in the NCBI PubChem database, enabling access to additional compound information.

In the “Variation” module, the database presents variations of resequencing data of 149 accessions (Table S3), including SNPs and InDels. Users first select the species and input gene ID to display corresponding results, and then can search by entering variation ID, chromosomes, variation types, and more. Based on the robust annotation capabilities and international recognition of ANNOVAR, we used it to annotate SNPs and indels detection results for various species (Table S4). We also provide basic information about samples from different projects, such as tissue, age, developmental stage, and locality. Furthermore, all data are available for users to download.

In addition to the main modules, the database contains 11 popular bioinformatics tools located in the navigation bar, including “Gene Search”, “Gene Function”, “Gene Correlation”, “Transcription Factor”, “BLAST”, “KEGG Enrichment”, “GO Enrichment”, “Primer3 web”, “Genome Synteny”, “Jbrowse” and “Expression Heatmap”. For instance, the BLAST tool is incorporated into a stand-alone web page, where 23 genome assemblies are available for querying orthologous gene candidates. The “Expression Heatmap” functionality offers an intuitive means of data visualization, aiding researchers in swiftly comprehending gene expression data and identifying potential biological patterns and differences. The “Genome Synteny” tool is available for detection and evolutionary analysis of gene synteny and collinearity between magnoliids plants. The “Gene Correlation” tool is used to identify genes that are highly correlated with a target gene. It provides data for 14 species, allowing users to input a gene of interest into the search box to search for and download the IDs of highly correlated genes along with their correlation coefficients. The “Primer3 web” is a tool to design primers. The “Jbrowse” tool provides a fast and interactive genome browser for navigating large-scale high-throughput sequencing data under a genomic framework, and 16 available MagnoliidsGDB genomes are incorporated into JBrowse. Furthermore, we built a “Download” web page to allow users to freely obtain all the data available in MagnoliidsGDB. These data include genome assembly sequences, annotation results, transcriptomic analyses results, and metabolomics data.

MagnoliidsGDB aims to provide a comprehensive multi-omics data and powerful tools to enable researchers obtained available information conveniently and smoothly. For instance, magnoliids can be classified into three types: vines (e.g., *Piper nigrum*), shrubs (e.g., *Chimonanthus salicifolius*), and trees (e.g., *Cinnamomum kanehirae*) based on their growth habits and structural characteristics. Vines usually need to cling to additional structures such as trees, fences, or rocks for support; shrubs typically lack a prominent main trunk, exhibiting a clustered growth form and relatively short stature. In contrast, trees possess a distinct main trunk and are generally tall, with some species reaching heights exceeding 100 meters. It is well-known that lignin plays crucial roles in providing structural support and facilitating water and nutrient transport. Therefore, we speculate that differences in lignin biosynthesis among these three plant types result in variations in lignin accumulation, leading to their different growth habits. To confirm this hypothesis, we identified homologous genes related to lignin biosynthesis in the aforementioned representative species using the BLAST tool in MagnoliidsGDB. Using the expression heatmap tool in the transcriptome module, we observed the real differences in the expression of these genes in the stem, leaf, and flower tissues between *C. kanehirae, C. salicifolius* and *P. nigrum* (Figure 1D), which were consistent with our speculation. Furthermore, we employed the genome synteny tool to generate a chromosomal localization map of these genes (Figure 1E), showing their genomic variations in magnollids.

In summary, we present MagnoliidsGDB, an authoritative and convenient platform providing valuable resources for the research of magnoliids. In the future, MagnoliidsGDB will continuously update newly published data, and add other omics data, such as proteomics, along with powerful tools for users. MagnoliidsGDB is expected to become a central community portal for magnoliids research and provide long-term support for the research of plant evolution.

## Acknowledgments

This work was supported by the National Natural Science Foundation of China (32160058). Special thanks to Hao Wang from BMW Brilliance Automotive Ltd. for his support on the implementation of the “Genome Synteny” tool’s construction of MagnoliidsGDB. We also extend our gratitude to Yechun Xu from Institute of Environmental Horticulture, Guangdong Academy of Agricultural Sciences, for his sharing of high-quality figures of many magnoliids in the MagnoliidsGDB.

## Author contributions

Yuanhao Ding and Zhuang Yang conceived the project. Yayu Chen, Jiajie Chen, Ping Li, and Xinglong Zhao performed the bioinformatics analysis. Yayu Chen collected, analysed, and visualized the omics data and developed the tools in MagnoliidsGDB. Yayu Chen, Sihui Huang, Zhumao Li, and Sishu Huang collected the samples, obtained, and proceeded the metabolic data. Yayu Chen and Yuanhao Ding wrote the manuscript. Yuanhao Ding, Zhuang Yang, Haiyan Hu, and Jie Luo discussed the study and revised the manuscript. All authors read and approved the final manuscript.

## Supplemental information

Table S1. Transcriptome data information of 20 species.

Table S2. Metabolome data information of 15 species.

Table S3. Resequencing data information of 6 species.

Table S4. Statistical summary of SNPs and Indels detection results.

